# The distribution of antibiotic use and its association with antibiotic resistance

**DOI:** 10.1101/473769

**Authors:** Scott W. Olesen, Michael L. Barnett, Derek R. MacFadden, John S. Brownstein, Sonia Hernández-Díaz, Marc Lipsitch, Yonatan H. Grad

## Abstract

Antibiotic use is a primary driver of antibiotic resistance. However, antibiotic use can be distributed in different ways in a population, and the association between the distribution of use and antibiotic resistance has not been explored. Here we tested the hypothesis that repeated use of antibiotics has a stronger association with population-wide antibiotic resistance than broadly-distributed, low-intensity use. First, we characterized the distribution of outpatient antibiotic use across US states, finding that antibiotic use is uneven and that repeated use of antibiotics makes up a minority of antibiotic use. Second, we compared antibiotic use with resistance for 72 pathogen-antibiotic combinations across states. Finally, having partitioned total use into extensive and intensive margins, we found that intense use had a weaker association with resistance than extensive use. If the use-resistance relationship is causal, these results suggest that reducing total use and selection intensity will require reducing broadly-distributed, low-intensity use.

## Introduction

Antibiotic use is a primary driver of antibiotic resistance, and reducing antibiotic use is a central strategy for combatting resistance (1, 2). Understanding the relationship between antibiotic use and antibiotic resistance is therefore critical for the design of rational antibiotic stewardship strategies. Multiple studies have identified cross-sectional relationships between antibiotic use and resistance, especially across European countries and US states (3–9). In general, these studies compare total outpatient antibiotic use with population-level resistance. However, antibiotic use is generally not evenly distributed. A study of outpatient prescribing in the UK found that 30% of patients were prescribed at least one antibiotic per year, with the top 9% of patients receiving 53% of all antibiotics (10). A study of beneficiaries of Medicare, a national health insurance program that covers that vast majority of Americans 65 and older, found that the proportion of beneficiaries who take antibiotics varies by US state and drug class (11). In some cases, antibiotic courses can last for months or even years (12, 13). Because antibiotic use is uneven, total use does not distinguish between broad use—many people receiving a few prescriptions—and intense use—a few people receiving many prescriptions (14).

It stands to reason that the distribution of antibiotic use, not just total use, could have an effect on resistance (15). There are a few studies of the relationship between repeated antibiotic exposure on antibiotic resistance (16–22), and it remains unclear whether broad use or intense use is associated with population-level resistance. For example, if a first course of antibiotics given to an antibiotic-naive patient clears most of the susceptible bacteria they carry, then a second course in the same patient will have only a small effect, since most susceptible bacteria were already eliminated. Giving that second course to a different, antibiotic-naive patient instead would have a greater effect on population-level resistance. On the other hand, multiple courses given to a single patient might have a synergistic effect on resistance.

The goal of this study was to test the hypothesis that intense antibiotic use has a stronger association with population-level resistance than broad, low-level antibiotic use. We used an ecological design to compare the distribution of antibiotic use with antibiotic resistance. Although an ecological design is potentially subject to confounders and cannot definitively test for the causal effect of the distribution of use on resistance at the individual level, ecological studies of use and resistance are the most feasible design for studying the relationship between antibiotic use and population-level resistance, and the results of ecological designs play an important role in developing antibiotic stewardship policies (23, 24).

To test this hypothesis, we first characterized the distribution of outpatient antibiotic use in two US nationwide pharmacy prescription claims databases, Truven Health MarketScan Research Database (25) and Medicare, both covering 2011-2014. We considered only outpatient antibiotic prescribing, which accounts for 80-90% of total medical antibiotic use in the UK and Sweden (26, 27) and is presumed to account for a similar fraction in the US (28). Unlike antibiotic sales data and nationwide healthcare surveys (29), MarketScan and Medicare claims data, which have previously been used to characterize variations in antibiotic use (11, 30–32), provide longitudinal prescribing information about individual people, which can distinguish between many people getting a few prescriptions and a few people getting many prescriptions. We characterized the distribution of antibiotic use across US states by partitioning annual total use as the sum of annual first use—individuals’ first pharmacy fill for an antibiotic in a calendar year—and annual repeat use—pharmacy fills beyond individuals’ first ones in a calendar year. Second, we compared annual total antibiotic use with antibiotic resistance as measured in ResistanceOpen, a US nationwide sample of antibiotic susceptibility reports, for 2012-2015 (i.e., lagged by one year (8, 33)), evaluating the relationship between use and resistance across US states for 72 pathogen-antibiotic combinations. Finally, we evaluated whether annual first use and annual repeat use are differently associated with population-level resistance.

## Results

### Antibiotic use is not evenly distributed

Our analysis included 99.8 million outpatient pharmacy antibiotic prescription fills among 62.4 million unique people, approximately 20% of the US population, during 2011-2014 using the MarketScan database (25). In 2011, 34% of people received an antibiotic, and 10% of people received 57% of all antibiotic prescriptions. This distribution varied by population but was similar across data years (Figure 1 - Figure Supplement 1). To characterize the distribution of specific antibiotics, we grouped individual antibiotic generic formulations into drug groups based on their chemical structures and mechanisms of action (Supplementary File 1 - Table 1). For all drug groups, most people had zero prescriptions for that antibiotic in a given year, but antibiotics differed in their distributions (Figure 1).

**Figure 1:**
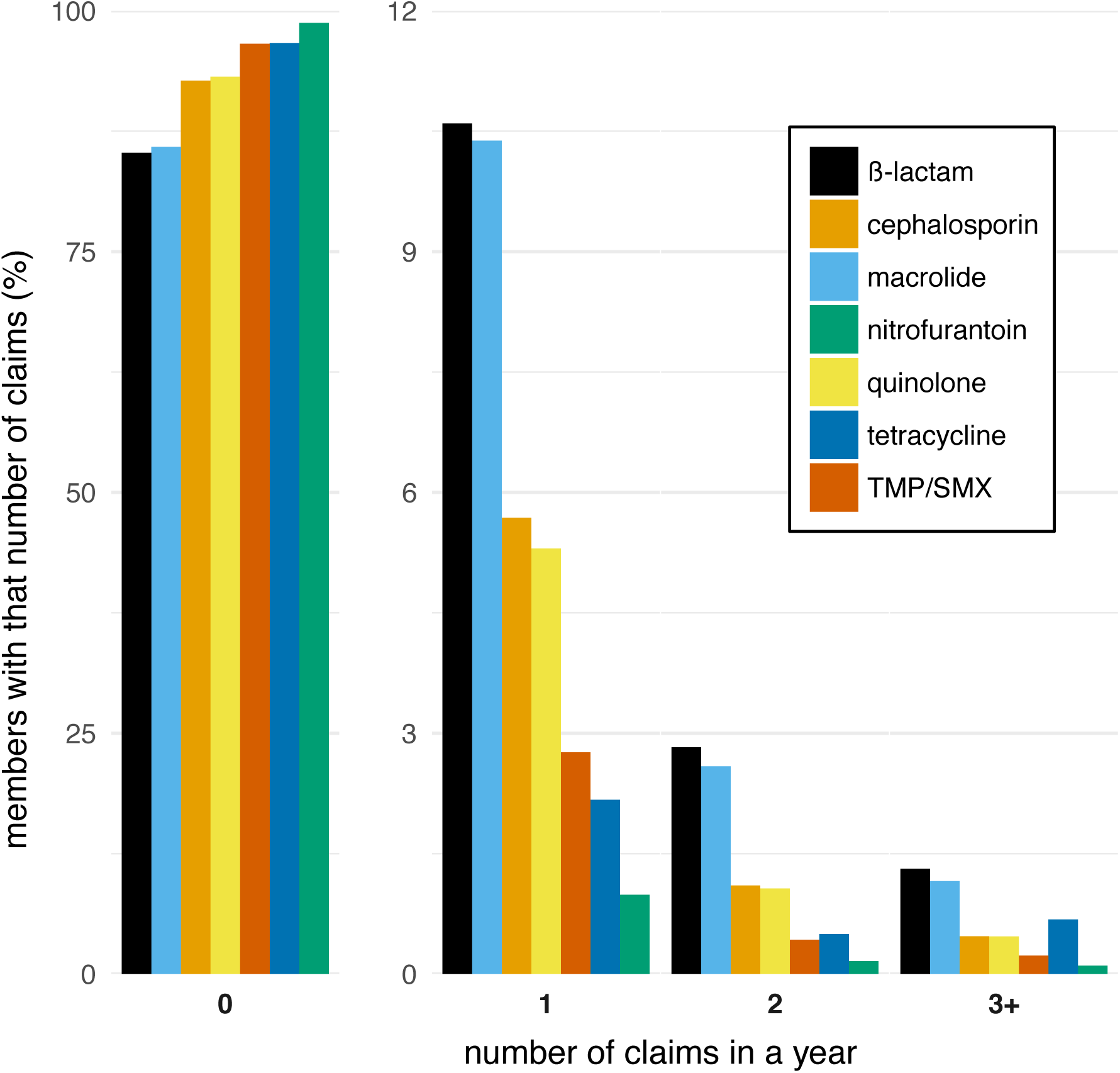
The distribution of antibiotic use within individuals. Bars indicate the proportion of members in the MarketScan data with different numbers of prescription fills in 2011 for each of the drug groups. TMP/SMX: trimethoprim/sulfamethoxazole.

We next examined the distribution of antibiotic use for each drug group and US state. To quantify the distribution of antibiotic use, we labeled each antibiotic pharmacy claim as “first” if it was the first pharmacy fill for that drug group made by that individual in that calendar year, and “repeat” if it was a second, third, etc. fill for an antibiotic in the same drug group made by the same individual in the same calendar year. An individual’s first and repeat claims in a calendar year add up to their total number of claims for that year. We then partitioned population-level annual total use, measured as pharmacy fills per 1,000 members per year, into the sum of annual first use, measured as first fills per 1,000 members per year, and repeat use, measured as repeat fill per 1,000 members per year, for each drug group and US state. Annual first use of a drug group is equivalent to the proportion of the population taking an antibiotic in that group in that year.

Total use varied between drug groups and across states (Figure 2). Annual repeat use made up a steady one-quarter to one-third of annual total use across drugs and states, with the exception of tetracyclines, for which high repeat use was associated with young adults (Figure 2 – Figure Supplements 1 and 2), probably for acne treatment. This distribution of first and repeat use is distinct from the pattern predicted by the single-parameter Poisson and geometric distributions (Figure 2), but the ratio of first use to repeat use for each drug was nearly constant across US Census regions (Figure 2 – Figure Supplement 3). Thus, the higher antibiotic use in the Southern states (11, 34) is primarily attributable to a greater proportion of people taking antibiotics, not because those who receive antibiotics receive more of them.

**Figure 2:**
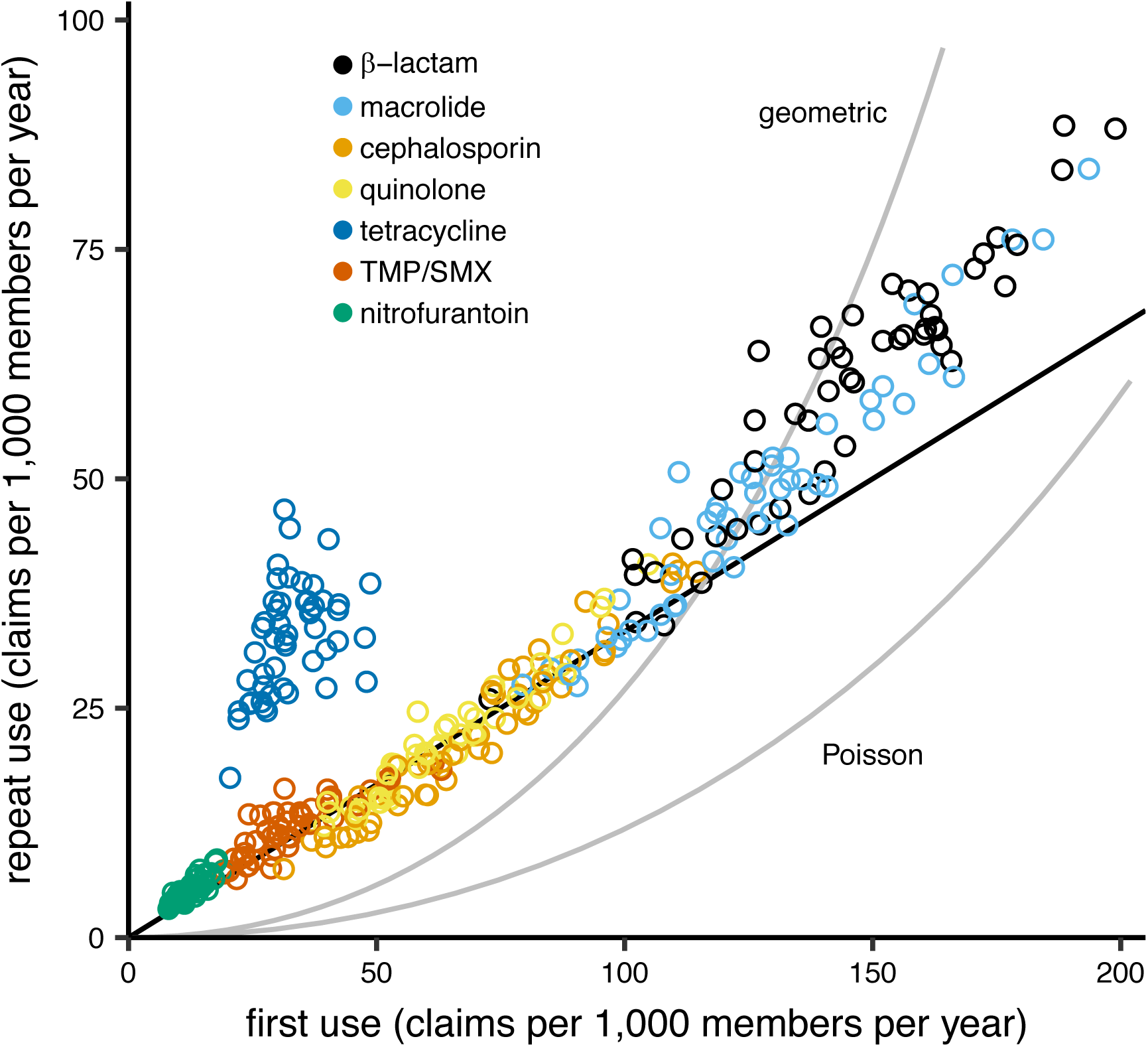
The distribution of antibiotic use across US states. Each point indicates first use and repeat use of a single drug group in a single US state (averaged over the data years). Points falling on the black line have three times as much first use as repeat use (i.e., repeat use is one- quarter of total use). The curves show the relationships between first use and repeat use expected from the Poisson and geometric distributions. TMP/SMX: trimethoprim/sulfamethoxazole.

### Landscape of correlations between total use and resistance across pathogens and antibiotics

To verify that our antibiotic use and resistance data sources could be used to distinguish the associations of first use and repeat use with antibiotic resistance, we first measured the landscape of Spearman correlations between total use and antibiotic resistance for multiple pathogens and antibiotics (3–8). To measure antibiotic resistance, we used a US nationwide sample of hospital antibiotic susceptibility reports (35), which included resistance of 38 pathogens to 37 antibiotics in 641 antibiotic susceptibility reports from 230 organizations (hospitals, laboratories, and surveillance units) spread over 44 US states. Although most organizations contributing antibiotic susceptibility reports were hospitals, hospital antibiotic susceptibility reports are biased toward community-acquired organisms (36, 37), and studies often compare hospital antibiotic susceptibility reports with community antibiotic use (38).

Because the epidemiology and pharmacology of each pathogen-antibiotic combination is unique, each combination could have a unique use-resistance relationship (15). We therefore aggregated antibiotic resistance into the same drug groups with which we aggregated antibiotic use (Supplementary File 1 – Table 1) and evaluated the 72 pathogen-antibiotic combinations that were adequately represented in the antibiotic resistance data (see Methods). Across those 72 combinations, correlation coefficients ranged from –32% to 64% (Figure 3, Supplementary File 1 - Table 2). The strongest correlation (Spearman’s *ρ* = 64%, 95% CI 41 to 80%) was between macrolide use and the proportion of *Streptococcus pneumoniae* isolates that were macrolide nonsusceptible (Figure 4). Correlation coefficients were mostly positive (median correlation coefficient 21%, IQR 8 to 34%). Use-resistance correlations involving macrolides, quinolones, and cephalosporins were more positive than those for nitrofurantoin, and correlations involving quinolones were more positive than those for trimethoprim/sulfamethoxazole (pairwise Mann-Whitney tests, two-tailed, FDR = 0.05). Coefficients were not significantly more positive for any particular pathogen.

**Figure 3:**
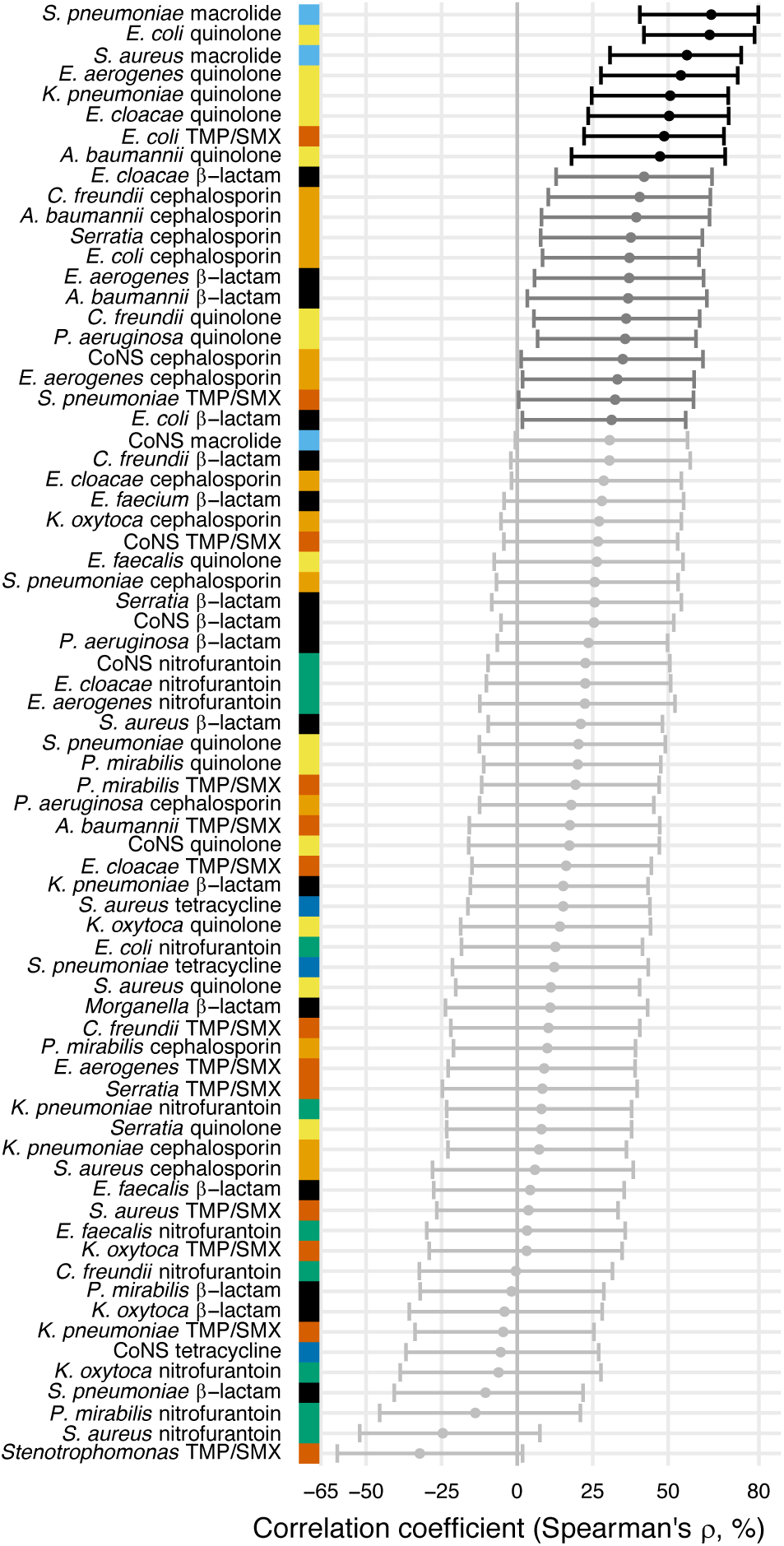
Correlations between total antibiotic use and resistance are biased toward positive values. Error bars show 95% confidence intervals. The color strip visually displays the drug groups. Statistical significance is indicated by color of the points (black, significant at FDR = 0.05, two-tailed; dark gray, significant at α = 0.05, two-tailed; light gray, not significant). TMP/SMX: trimethoprim/sulfamethoxazole. CoNS: coagulase-negative *Staphylococcus*.

**Figure 4:**
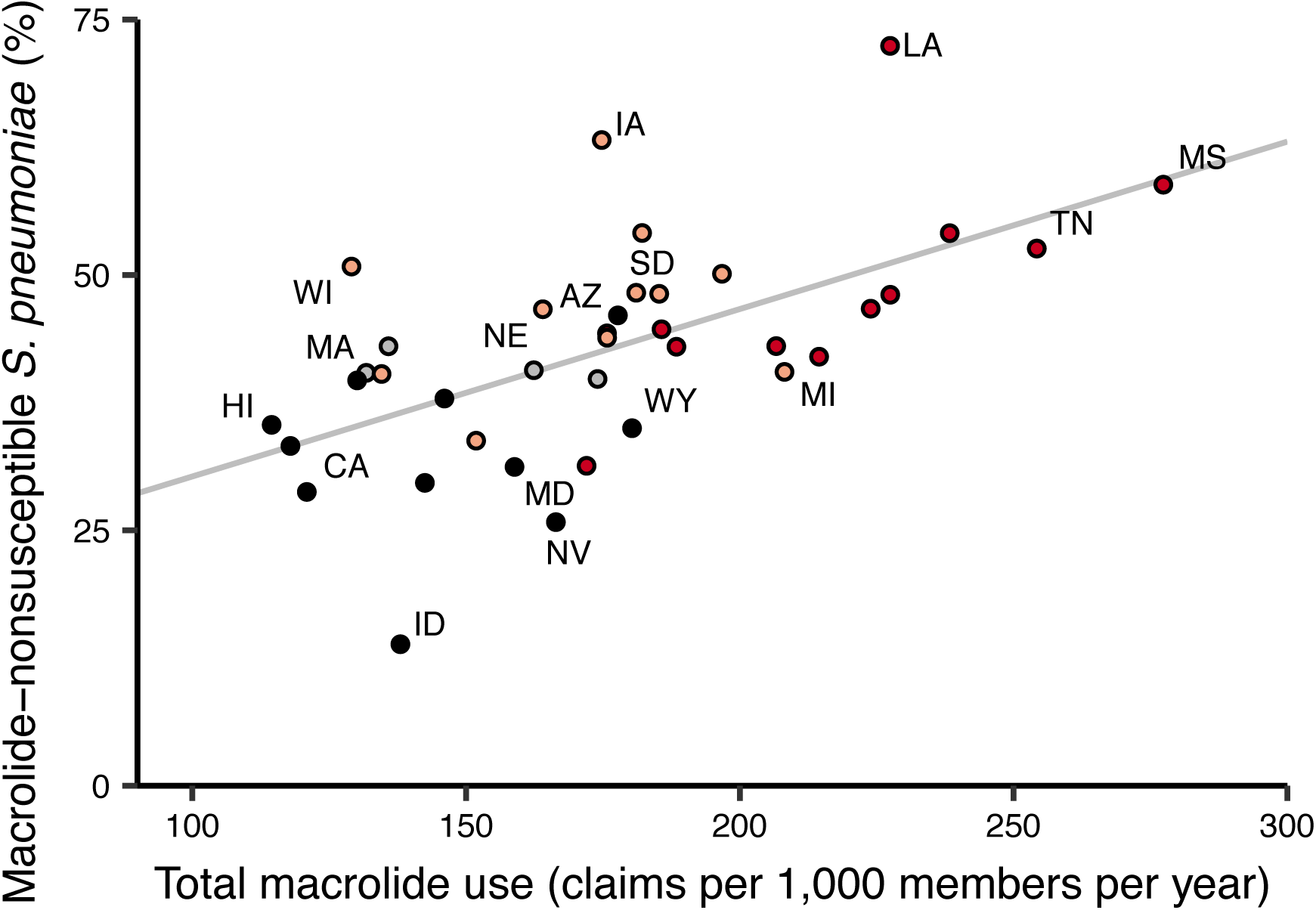
Total macrolide use and macrolide resistance among *Streptococcus pneumoniae* correlate across US states. Labels indicate selected states. Colors indicate US Census region (red, South; light red, Midwest; gray, Northeast; black, West). Line shows unweighted linear best fit. Southern states have highest macrolide use and resistance.

Because isolates from older adults are disproportionately represented in antibiotic susceptibility reports (39), we suspected that population-wide resistance might, in some cases, correlate better with antibiotic use among older adults. We therefore queried outpatient pharmacy antibiotic claims records from individuals 65 and older on Medicare (see Methods). When antibiotic use among Medicare beneficiaries was substituted for antibiotic use as measured in the MarketScan data (Figure 2 – Figure Supplement 2), correlation coefficients were similar (Supplementary File 1 – Tables 2 and 3). Conversely, children are the primary carriers for some pathogens (e.g., *Streptococcus pneumoniae* (40)), so we suspected that resistance might, in other cases, better correlate with children’s antibiotic use. Restricting the antibiotic use data to members at most 15 years old (Figure 2 – Figure Supplement 2) again yielded similar coefficients (Supplementary File - Tables 2 and 3). Thus, the landscape of correlations we observed was mostly robust to the exact population and data source.

We also evaluated the sensitivity of the results to the measurement of antibiotic use, substituting days supply of antibiotic for number of pharmacy fills, and the geographic level of the analysis, by aggregating the Medicare use data and resistance data at the level of the 306 hospital referral regions intended to approximate regional health care markets (41) (Supplementary File – Tables 2 and 3). The absolute values of the correlation coefficients were slightly closer to zero when using days supply rather than fills (Wilcoxon test, two-tailed; pseudomedian difference in absolute correlation coefficient 1.9 percentage points, 95% CI 0.72 to 3.1) and substantially closer to zero when using hospital referral regions rather than states as the units of analysis (6.1 percentage points, 95% CI 3.0 to 9.1).

### Lack of evidence for more positive association with repeat use

Having examined the landscape of the relationships between total use and resistance across pathogen-antibiotic combinations, we set out to test the hypothesis that repeat use has a stronger association with resistance than first use. For each pathogen-antibiotic combination, we performed a multiple regression predicting proportion nonsusceptible from first use and repeat use (Figure 5). First use and repeat use are highly correlated in some cases (Supplementary File -- Table 4) which will widen the confidence intervals on the regression coefficients but should not introduce bias (42). Regression coefficients for first use were more often positive than negative (54 of 72 [75%]; binomial test, 95% CI 63 to 84%). That is, first use was positively associated with resistance when controlling for repeat use. In contrast, regression coefficients for repeat use were more often negative than positive (44 of 72 [61%]; binomial test, 95% CI 49 to 72%). That is, repeat use was negatively associated with resistance when controlling for first use.

**Figure 5:**
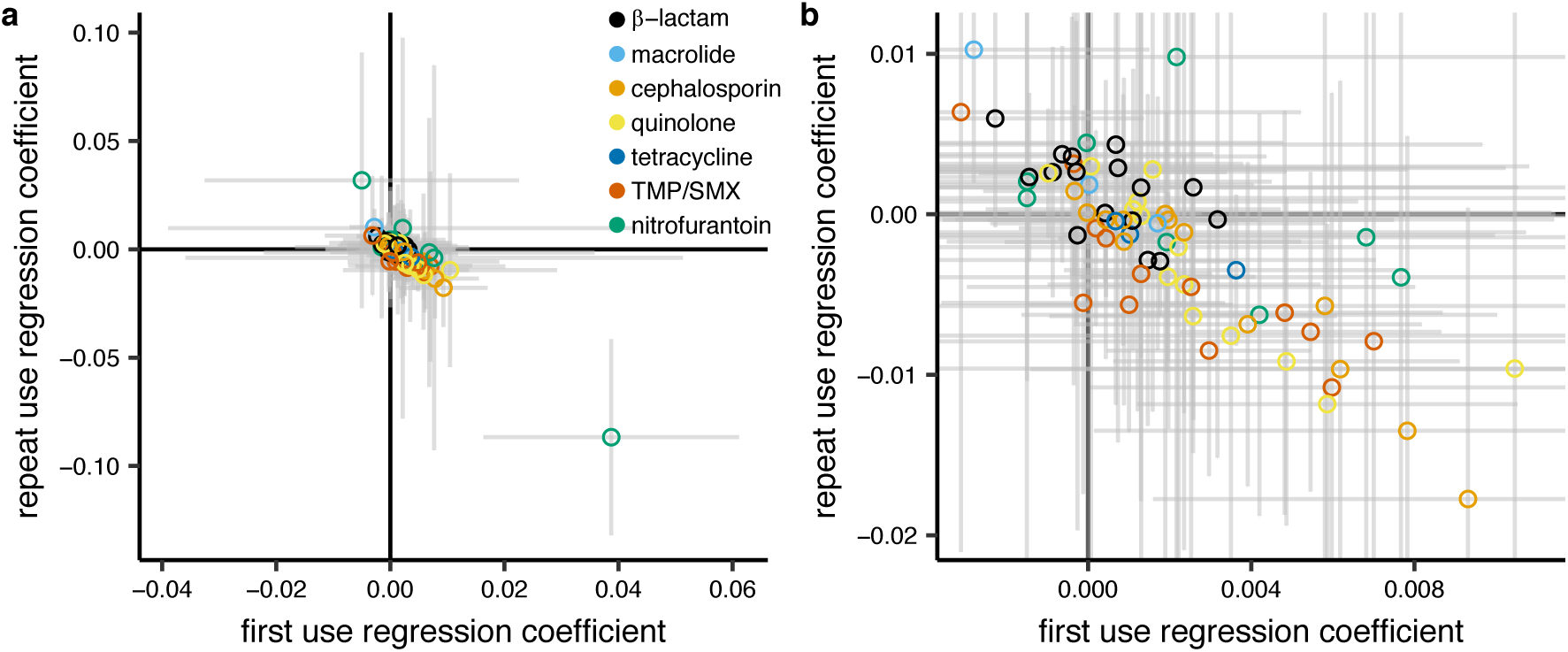
Repeat use tends to be negatively associated with resistance when controlling for first use. Each point represents a pathogen-antibiotic combination. The position of the point shows the two coefficients from the multiple regression. The units of the coefficients are proportion resistant per annual claim per 1,000 people. Color indicates drug group. Error bars show 95% CIs. (*a*) All data. (*b*) Same data, showing only the center cluster of points.

We evaluated the sensitivity of this result to age group, data source, metric of antibiotic use, and geographic unit of analysis, as described above. In all cases, regression coefficients for first use in the multiple regression were more likely to be positive than negative, while regression coefficients for repeat use were more likely to be negative than positive (Supplementary File 1 - Table 5). For certain pathogens and antibiotics, resistance could presumably accumulate in an individual over many years (18, 21), so we also computed alternate measures of first and repeat use by considering only individuals who were included in the MarketScan data for each year of 2011-2014, and we labeled an antibiotic fill as first use only if it was the first fill for that drug group made by that individual in the entire four-year period. In that analysis, a similar proportion of regression coefficients for first use were positive (69%, 95% CI 57 to 80%) and regression coefficients for repeat use were equally likely to be positive or negative (53%, 95% CI 41 to 65%).

## Discussion

### Landscape of use-resistance relationships

We used US nationwide datasets measuring antibiotic use in 60 million individual and antibiotic resistance in 3 million bacterial isolates to analyze relationships between antibiotic use and resistance, examining 72 pathogen-antibiotic combinations simultaneously, using identical data sources and analytical methods across combinations. Although previous studies have examined multiple pathogen-antibiotic combinations, usually no more than 5 pathogens or antibiotics are considered at once (3, 4, 8). We found that correlations between total use and resistance were mostly positive, that certain drugs tended to have more positive correlations, but that there was no clear pattern by organism (6). The overall landscape of correlations was mostly robust to the age groups studied and the geographic scale of the analysis, although correlations were somewhat weaker when conducting analysis at smaller geographic scales (8, 43–45). We used outpatient antibiotic use as the predictor of resistance because 80-90% of antibiotic use occurs in the outpatient setting (28) and because most antibiotic pressure on pathogens is due to “bystander selection”, in which the patient is treated for some reason other than an infection caused by that pathogen (46).

The correlations we observed between total antibiotic use and population-wide antibiotic resistance were noticeably weaker than those in highly cited European studies but comparable to those from other analyses of European data. For example, for *S. pneumoniae* and macrolides, Goossens *et al*. (3) reported a Spearman’s *ρ* of 83% and García-Rey *et al.* (4) reported 85%, while we found 62% and van de Sande-Bruinsma *et al*. (8) reported a median of 56%. These studies all used similar statistical methods, so the differences in the results must be due to some other factors (e.g., data quality, range of antibiotic use, distribution of antibiotic use, or pathogen biology). Correlations between *E. coli* resistance and use of β-lactams, cephalosporins, trimethoprim/sulfamethoxazole, and quinolones were similar to those reported in studies of use and resistance in UK primary care groups (43, 44).

The most notable difference, however, between our study and previous results from Europe is for *S. pneumoniae* and β-lactams: Goossens *et al*. (3) report a Spearman’s *ρ* of 84%, but we found no relation (–11%, 95% CI –41 to 22%). We propose that the narrow variation in β-lactam use across US states, approximately two-fold between the highest- and lowest-using states, obscures a correlation that is more apparent in Europe, where there is a four-fold variation between the highest- and lowest-using countries (3). Thus, our results and those from Goossens *et al*. may be consistent with respect to the underlying biology. We also note that, when reproducing the methodology from a US study (47) of the use-resistance relationship for β-lactams and *S. pneumoniae* (dichotomizing states as high- or low-prescribing and computing the odds ratio of resistance), we find a consistent point estimate but with wider confidence intervals (1.15, 95% CI 0.75 to 1.76).

Our study design may limit the interpretability of the landscape of use-resistance relationships. First, like the leading European studies using EARS-Net and US studies using the Centers for Disease Control and Prevention Active Bacterial Core surveillance, we compare population-wide outpatient antibiotic use with antibiotic susceptibility reports from hospitals. The degree to which hospital antibiotic susceptibility reports represent community infections is debated (36, 38). For example, if outpatient antibiotic use selects for resistance among community-acquired infections, and hospital antibiograms reflect data from community-acquired infections as well as unrelated inpatient resistance patterns, then the correlations we measure would be biased toward weaker associations. Furthermore, antibiotic use and resistance in the community setting is not completely independent of use and resistance in the hospital setting (48), and our approach does not account for any relationship between the two.

Second, antibiotic resistance is temporally dynamic, and our cross-sectional approach assumes that antibiotic use is autocorrelated across years (49) or resistance changes slowly (50). If use does cause resistance, and use and resistance changed meaningfully over the course of the study, then the correlations we measured by aggregating over all years would be biased toward weaker associations.

Third, because of the limitations in statistical power, we did not address the possibility that use of one antibiotic can select for resistance to another antibiotic (51, 52). Notably, use of one antibiotic can select for resistance to another antibiotic if the dominant clones of that species are resistant to both (51). In that case, if the use rates of the two drugs are correlated across states, then the apparent relationship between one drug and resistance to that drug would be biased upward. Furthermore, because the palette of antibiotic use varies by country (53), and different pathogen strains circulate in different populations, the univariate associations we observed between use of an antibiotic and resistance to that antibiotic in the US may not be applicable in other geographies.

Finally, like in other studies of antibiotic use, we did not address patient adherence, and typical approaches to address adherence using claims data (54) are problematic when the intended duration of treatment is not clear. The measured correlation would then be biased if, for example, poor patient adherence increased resistance and patient adherence were correlated with antibiotic use.

### Distribution of antibiotic use and antibiotic resistance

We described the distribution of antibiotic use across drug groups and US states, finding that 34% of the study population took an antibiotic in a year, and 10% of the population had 57% of the antibiotic fills in that year, similar to results from the UK (10), although this distribution varied by population (Figure 1 – Figure Supplement 1). By partitioning annual total use into annual first use and annual repeat use, we were able to show that, for each drug, annual first use makes up the majority of annual total use and that variations in annual first use explain more variance in annual total use than do variations in annual repeat use. We also found that first use tends to have a positive association with resistance when controlling for repeat use, while repeat use tends to have negative or associations with resistance when controlling for first use. This result held across sensitivity analyses.

If these associations are causal, that is, if outpatient first and repeat antibiotic use select for resistance among community-acquired pathogens, then our results would imply that antibiotic resistance in the outpatient setting is due more to first use, which tended to have positive associations with resistance, than to repeat use. In contrast to proposals to focus on intense antibiotic users for combatting resistance (10), this situation would imply that preventing marginal prescriptions among patients whose indications are borderline-appropriate or inappropriate for antibiotics may be the more effective tactic for reducing the prevalence of resistance mechanisms already established in the US.

There are limitations to the interpretability of these results. First, as mentioned above, there is a potential mismatch between the sources of the antibiotic use and antibiotic resistance data.

Second, although antibiotic use is a major driver of antibiotic resistance, the observed results may not be causal. Factors beyond antibiotic use, like population density, play a role in antibiotic resistance (2, 9). Even if antibiotic use and resistance are causally related, it may be that resistance affects antibiotic use. For example, if resistance to a drug is high, treatment using that drug is more likely to fail, discouraging repeated use, so that high resistance lead to decreased repeat use (36, 43, 51). Ecological studies like this one do not directly address causality, and further work is needed to distinguish between different causal pathways.

Third, the observed population-level relationships between antibiotic use and resistance need not also hold for the relationship between an individual’s first and repeat antibiotic use and the risk of a resistant infection in that individual. Any comparisons between our population-level results and individual-level studies would need to account for the difference between our population-level measures of first and repeat antibiotic use and the individual-level timing of antibiotic use and measurements of resistance.

Fourth, controlling for factors beyond antibiotic use could alter the apparent relationship between antibiotic use and resistance. In particular, we speculate that controlling for patient morbidity, which we did not address in this population-level analysis, would amplify the observed result, that first use tends to have a more positive association with antibiotic resistance than repeat use. We expect that morbid individuals have more repeat antibiotic use. We also expect that morbid individuals visit the hospital more often, putting them at higher risk of antibiotic resistant infections regardless of their antibiotic use. Thus, we speculate that repeat use causes resistance and also is a predictor of morbidity, which is associated with resistance. Failing to control for morbidity thus biases the association between repeat use and resistance toward more positive values. Conversely, controlling for morbidity would decrease the measured relationships between repeat use and resistance, amplifying our central result.

Fifth, we defined first and repeat use with respect to the calendar year, while it may be that some other timescale is the appropriate one for this analysis. Although our central result held when re- defining first and repeat use with respect to a four-year period (Supplementary File 1 - Table 5), it may be that, say, repeat use within an individual on a time-scale shorter than a year is an important determinant for risk of resistance in that individual. Our study does not distinguish between repeat use that occurs across year boundaries, which is presumably important for relating individuals’ antibiotic use with their risk of resistance.

Finally, we note that first use and repeat use are only one set of many ways of measuring the distribution of antibiotic use. For example, 10 repeat uses could mean 1 person with 10 repeat uses or 10 people with 1 repeat use each. The first and repeat use metrics cannot distinguish between these two cases, and it may be that some other measure of the distribution of antibiotic use would yield different results.

In conclusion, we find that population-wide antibiotic use and population-wide resistance appears to be more closely linked with broadly-distributed, low-intensity use rather than with intensity of use. Ultimately, accurate models predicting the emergence and spread of antibiotic resistance will require more careful characterizations of who gets what antibiotic (55), what selection pressure that places on pathogens, how those pathogens are transmitted, and in whom they manifest as infections (56). An ideal study would compare the complete history of an individual’s outpatient and inpatient antibiotic exposure with clinical microbiology data from that same individual, cross-referenced against population-level factors, among a representative, nationwide sample of individuals. Individual-level results could then also be compared with mechanistic models of resistance to draw inferences about within-host effects of antibiotic use (57, 58), and the role of co-occurring resistance and correlated antibiotic use could be addressed. In the absence of such a dataset, these ecological, associative results provide a guide to the development of antibiotic stewardship policy.

## Methods

### Study population and antibiotic use

MarketScan (25) data covering 2011-2014 were used to identify insurance plan members and characterize their outpatient antibiotic use. To ensure quality of the antibiotic use distribution data, only members who were on their insurance plan for 12 months during a given year were included. Prescription fills for oral and injected antibiotics were identified by generic formulation (Supplementary File 1 - Table 6) and drug forms (Supplementary File 1 - Table 7). We treated multiple fills on the same day for the same generic formulation with the same refill code as a single prescription fill. In the main analysis, antibiotic use was measured using fills, rather than days supply of drug, because some previous research has suggested that prescriptions better correlate with resistance (33) and that this choice is probably not detrimental (8, 49). The specific generic drugs were grouped into antibiotic drug groups designed to correspond to the antibiotic resistance drug groups described below (Supplementary File 1 - Table 1). All measures of antibiotic use were computed for each year 2011 to 2014, and the mean for each value across the 4 years was reported and used in analyses of the use-resistance relationship.

Antibiotic use among Medicare beneficiaries was measured as previously described (59). Briefly, we considered fee-for-service beneficiaries at least 65 years old among and with 12 months of enrollment in Medicare Part B and Part D among a 20% sample of beneficiaries for each of 2011-2014. The Medicare data, which provides the zip code for each beneficiary, were also aggregated at the level of hospital referral region (41), using the 2011 zip code to region crosswalk. MarketScan data do not include zip code-level resolution.

This study was deemed exempt from review by the institutional review board at the Harvard T. H. Chan School of Public Health.

### Antibiotic resistance

Antibiotic resistance prevalences for common bacterial pathogens were identified from ResistanceOpen, a previously developed database of spatially localized patterns of antibiotic resistance (35). This continuously updated database contains antibiotic resistance data from online sources during 2012 to 2015. At the time of analysis, the resistance data consisted of approximately 86,000 records, each indicating the fraction of isolates of an organism that were nonsusceptible to a particular drug in a particular antibiotic susceptibility report (“antibiogram”). The median number of isolates corresponding to each record was 93, but records had up to 75,000 associated isolates. 7 records (<0.01%) with missing numbers of isolates were excluded. In antibiograms that separated *S. aureus* into MRSA and MSSA, resistance of aggregate *S. aureus* to individual drugs was taken as the average of the MRSA and MSSA records, weighted by number of isolates. MRSA and MSSA were not considered as separate species in any analysis.

The specific antibiotics used in antibiotic resistance assays were grouped into antibiotic drug groups (Supplementary File 1 - Table 1) designed to correspond to the antibiotic use groups. If resistance to more than one antibiotic in a drug group was reported for a particular pathogen in a particular antibiogram, resistance to that drug group for that pathogen in that antibiogram was computed as the mean of the resistances measured for the antibiotics in that group, weighted by the number of isolates. The proportion of nonsusceptible isolates in a state for a particular pathogen-antibiotic combination was computed as the average of the proportions from each contributing antibiogram in that state, weighted by number of isolates.

### Statistical methods

Antibiotic use and resistance were compared using Spearman correlations and multiple linear regressions. Of the 887 pathogen-antibiotic combinations present in the data, we analyzed the 72 combinations that were present in at least 34 states. This excluded 21% of the pathogen-antibiotic-antibiogram combinations. We established the cut-off for number of states because 80% power to detect a Pearson correlation coefficient of magnitude 0.55 at *α* = 0.01 under a two-sided hypothesis requires at least 34 samples. (Although we report Spearman correlations, there is no straightforward power calculation methodology for Spearman correlations, and we used the Pearson power calculation as an approximation.) We aggregated data across all years, rather than comparing use and resistance in each year, because of the sparse of the resistance data: of 2,767 pathogen-antibiotic-state combinations in the data, only 182 have data for all four years. No pathogen-antibiotic combination had more than 4 states with data for all 4 years. Confidence intervals on correlation coefficients were computed using the Fisher transformation and normal approximation method. Multiple comparisons were accounted for using the Benjamini-Hochberg false discovery rate (FDR) (60). Multiple regressions predicted proportion of isolates nonsusceptible from first use and repeat use.

### Data availability

State-level, aggregate antibiotic use and resistance data used in the main analyses are in Figure 3 -Source data 1 and 2. We do not own and cannot publish disaggregated MarketScan or Medicare data. MarketScan data are available by commercial license from Truven Health (marketscan.truvenhealth.com). Medicare data are available from ResDAC (www.resdac.org). ResDAC requires an application ensuring that requesting researchers comply with Common Rule, HIPAA, and CMS security and privacy requirements. Disaggregated ResistanceOpen data are restricted due to hospitals’ privacy concerns. ResistanceOpen data are available by request from HealthMap (www.resistanceopen.org).

## Acknowledgements

SWO and ML were supported by cooperative agreement U54GM088558 from the National Institute of General Medical Sciences. The content is solely the responsibility of the authors and does not necessarily represent the official views of the National Institute of General Medical Sciences or the National Institutes of Health.

---

**Figure 1 - Figure Supplement 1.**
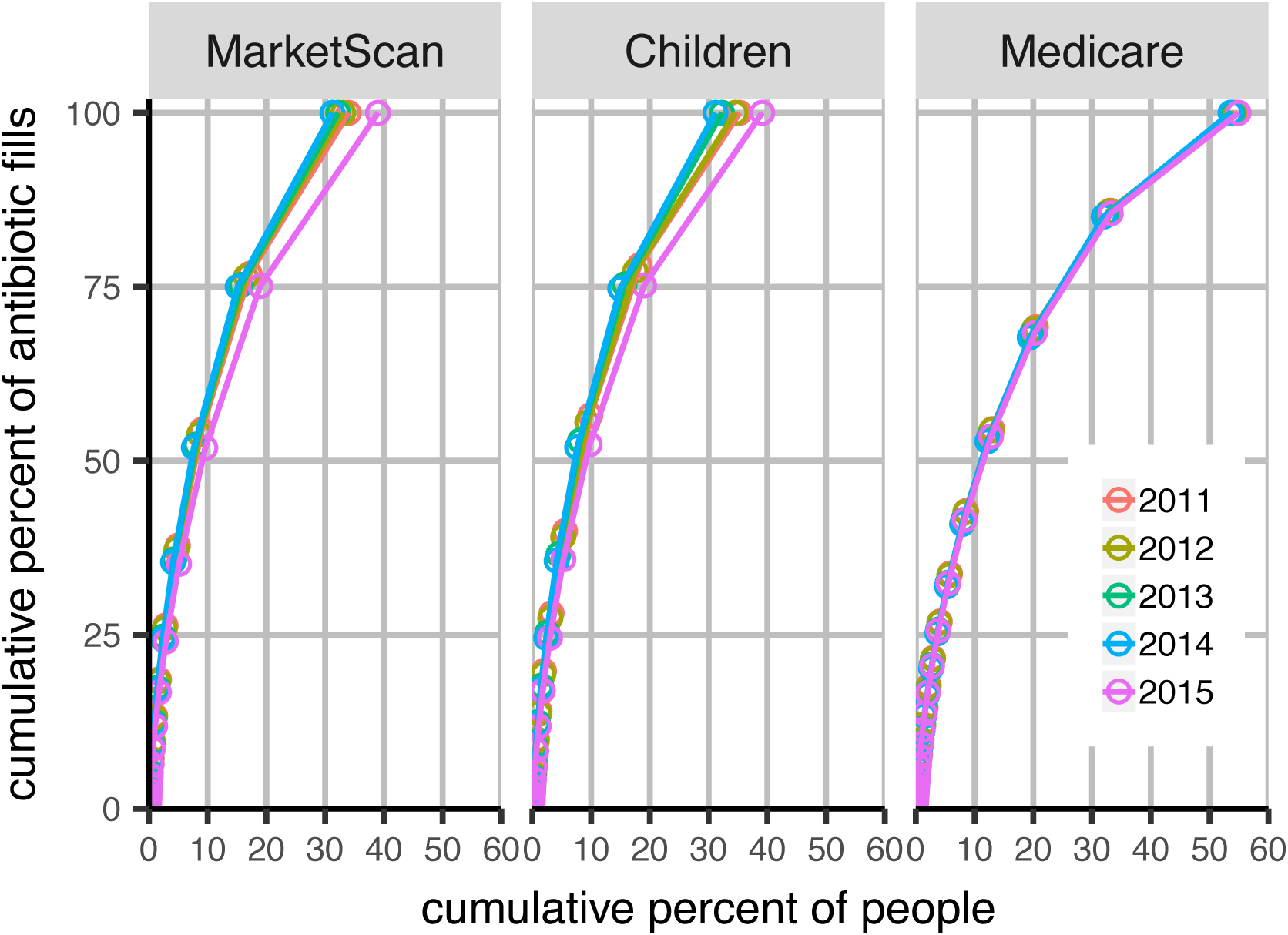
Cumulative distribution of antibiotic use. Each point represents a group of people with a certain number of associated claims for any antibiotic, starting at the left with the members with the greatest number of claims. The upper-right line segment shows members with 1 claim, the next segment shows members with 2 claims, etc. Colors indicate data years. Panels indicate study population. MarketScan: main data set. Children: MarketScan data including only members 15 and younger.

**Figure 2 – Figure Supplement 1.**
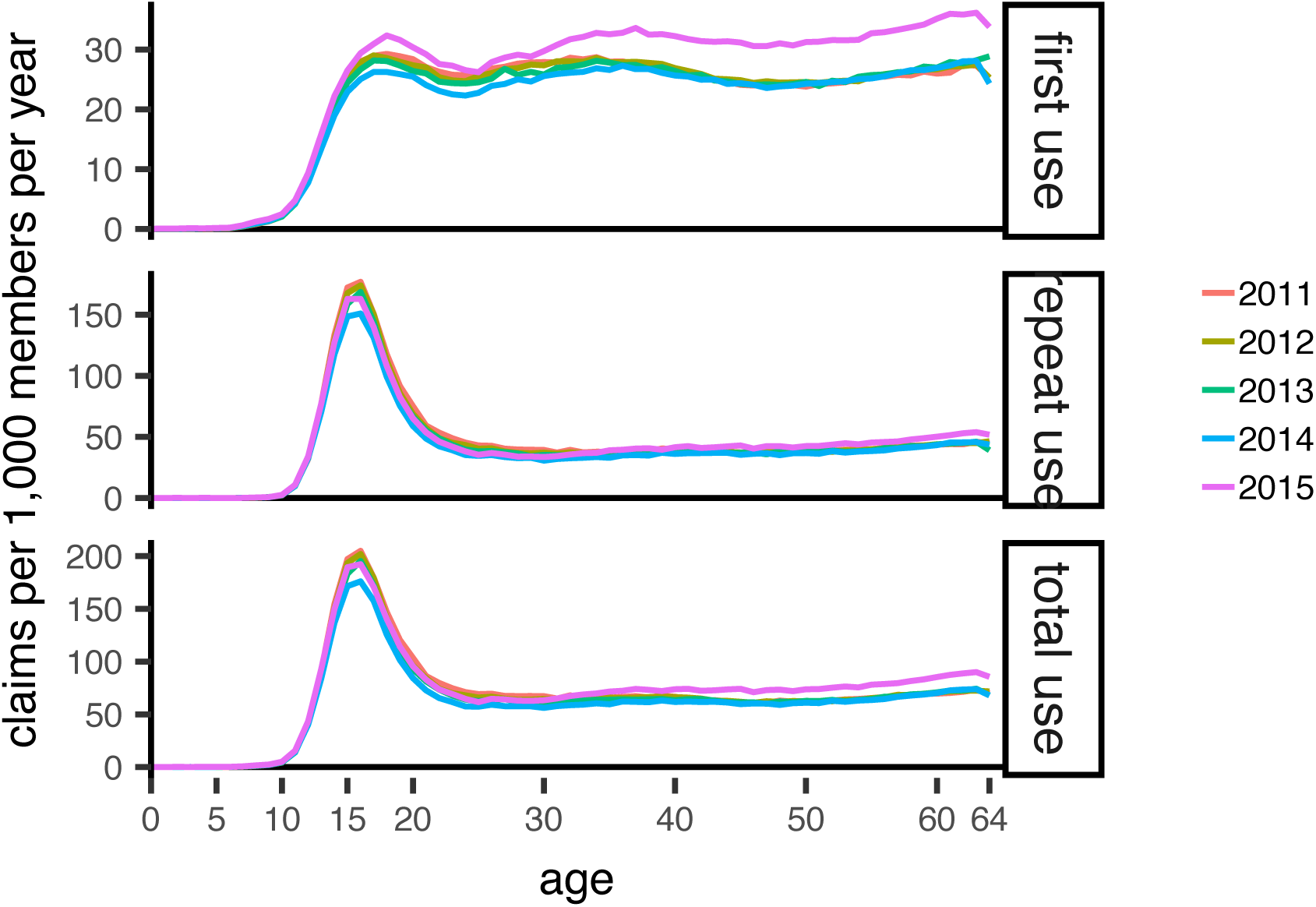
Distribution of tetracycline use by age. Colors indicate data years.

**Figure 2 – Figure Supplement 2.**
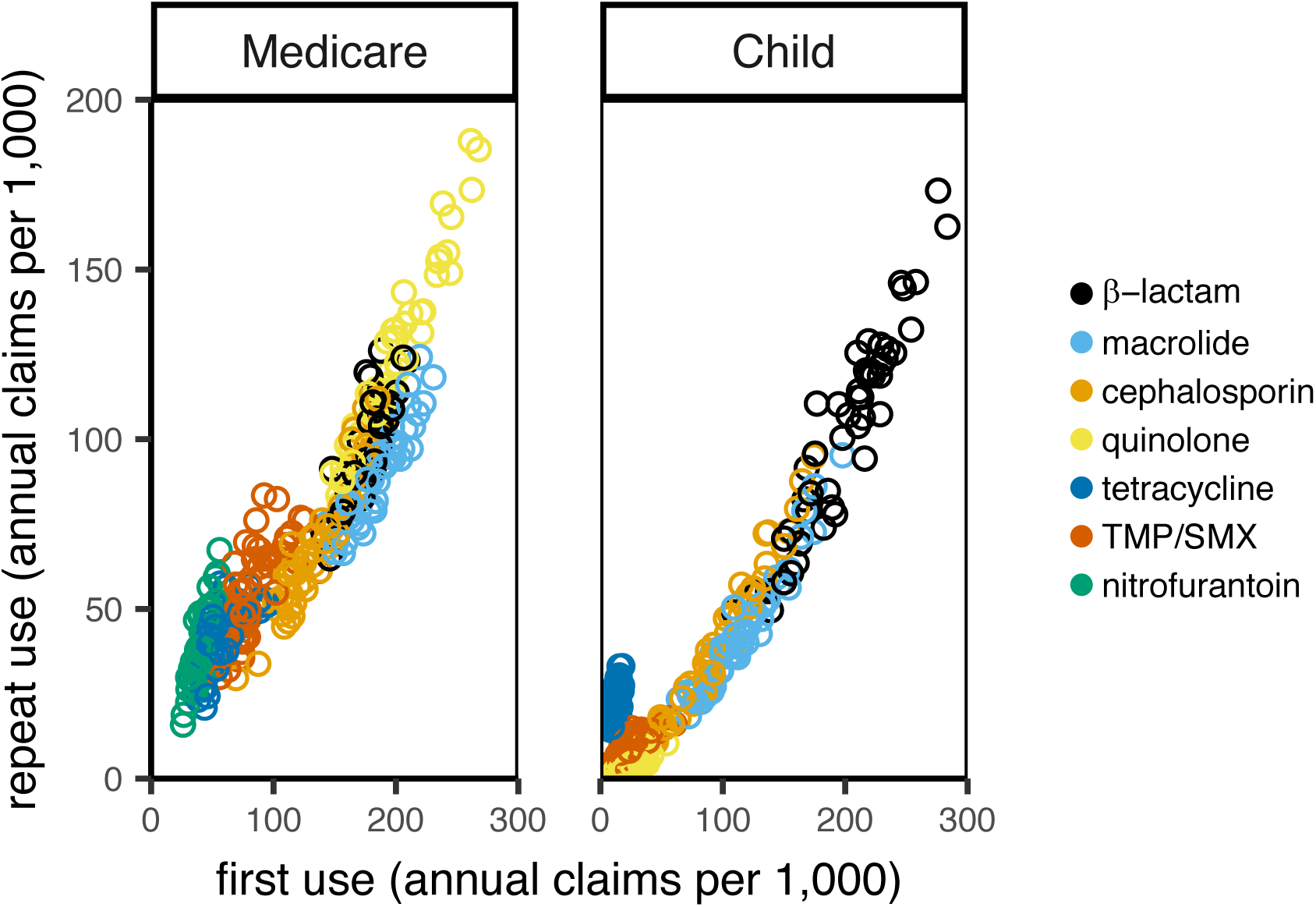
Distribution of antibiotic use by population. Each point represents average use of a drug group in a state across data years. Children: MarketScan data including only members 15 and younger.

**Figure 2 – Figure Supplement 3.**
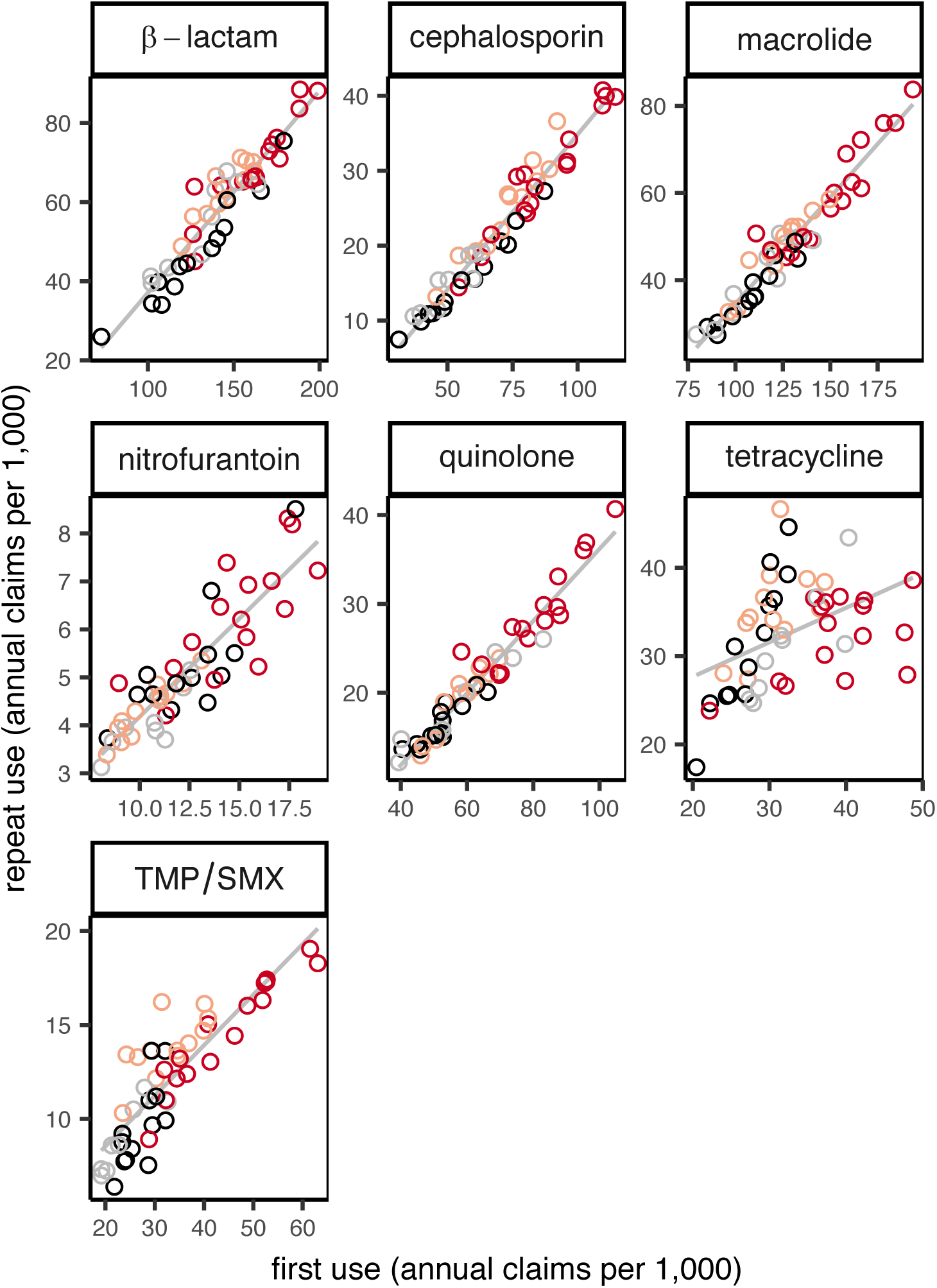
Distribution of antibiotic use by region. Each point shows use for a drug group in a state, averaged over data years. Colors indicate US Census region (red, South; light red, Midwest; gray, Northeast; black, West). Line shows unweighted linear best fit.

**Supplementary File 1.** Supplemental tables.

Figure 3 – Source data 1. **Antibiotic use data.** For each data source (MarketScan or Medicare), data subset or population among MarketScan records, state (with masked ID), and drug group, annual first and repeat claims per 1,000 members. Main: main data set. Children: members at most 15 years old. Days supply: first and repeat use are reported as days supply, not claims. Multiyear: among members in the data for all 4 data years.

Figure 3 – Source data 2. **Antibiotic resistance data.** For each adequately-represented pathogen and drug group (see Methods) and state (with masked ID matching the antibiotic use data), the proportion of isolates collected in that state susceptible to that drug.

## References

1. Gould, M I (1999) A review of the role of antibiotic policies in the control of antibiotic resistance. J Antimicrob Chemother 43(4):459–465.

2. Harbarth S, Samore MH (2005) Antimicrobial resistance determinants and future control. Emerg Infect Dis 11(6):794–801.

3. Goossens H, Ferech M, Vander Stichele R, Elseviers M (2005) Outpatient antibiotic use in Europe and association with resistance: a cross-national database study. The Lancet 365(9459):579–587.

4. Garcia-Rey C, Aguilar L, Baquero F, Casal J, Dal-Re R (2002) Importance of local variations in antibiotic consumption and geographical differences of erythromycin and penicillin resistance in Streptococcus pneumoniae. J Clin Microbiol 40:159–64.

5. Bronzwaer SLAM, et al. (2002) A European study on the relationship between antimicrobial use and antimicrobial resistance. Emerg Infect Dis 8(3):278–282.

6. Bell BG, Schellevis F, Stobberingh E, Goossens H, Pringle M (2014) A systematic review and meta-analysis of the effects of antibiotic consumption on antibiotic resistance. BMC Infect Dis 14:13.

7. European Centre for Disease Prevention and Control, European Food Safety Authority, European Medicines Agency (2017) ECDC/EFSA/EMA second joint report on the integrated analysis of the consumption of antimicrobial agents and occurrence of antimicrobial resistance in bacteria from humans and food‐producing animals. EFSA J 15. doi:10.2903/j.efsa.2017.4872.

8. van de Sande-Bruinsma N, et al. (2008) Antimicrobial drug use and resistance in Europe. Emerg Infect Dis 14(11):1722–1730.

9. MacFadden DR, McGough SF, Fisman D, Santillana M, Brownstein JS (2018) Antibiotic resistance increases with local temperature. Nat Clim Change 8(6):510–514.

10. Shallcross L, Beckley N, Rait G, Hayward A, Petersen I (2017) Antibiotic prescribing frequency amongst patients in primary care: a cohort study using electronic health records. J Antimicrob Chemother 72(6):1818–1824.

11. Zhang Y, Steinman MA, Kaplan CM (2012) Geographic variation in outpatient antibiotic prescribing among older adults. Arch Intern Med 172:1465–1471.

12. Enzler MJ, Berbari E, Osmon DR (2011) Antimicrobial prophylaxis in adults. Mayo Clin Proc 86:686–701.

13. Lau JSY, et al. (2017) Surveillance of life-long antibiotics: a review of antibiotic prescribing practices in an Australian Healthcare Network. Ann Clin Microbiol Antimicrob 16(1):3.

14. Berrington A (2010) Antimicrobial prescribing in hospitals: be careful what you measure. J Antimicrob Chemoth 65:163–168.

15. Turnidge J, Christiansen K (2005) Antibiotic use and resistance--proving the obvious. Lancet Lond Engl 365(9459):548–549.

16. Costelloe C, Metcalfe C, Lovering A, Mant D, Hay AD (2010) Effect of antibiotic prescribing in primary care on antimicrobial resistance in individual patients: systematic review and meta-analysis. BMJ 340:c2096.

17. Hillier S, et al. (2007) Prior antibiotics and risk of antibiotic-resistant community-acquired urinary tract infection: a case-control study. J Antimicrob Chemother 60:92–9.

18. Carothers JJ, et al. (2007) The relationship between previous fluoroquinolone use and levofloxacin resistance in Helicobacter pylori infection. Clin Infect Dis 44:e5–8.

19. Arason VA, et al. (1996) Do antimicrobials increase the carriage rate of penicillin resistant pneumococci in children? Cross sectional prevalence study. BMJ 313(7054):387–391.

20. Nasrin D, et al. (2002) Effect of beta lactam antibiotic use in children on pneumococcal resistance to penicillin: prospective cohort study. BMJ 324:28–30.

21. McMahon BJ, et al. (2003) The Relationship among Previous Antimicrobial Use, Antimicrobial Resistance, and Treatment Outcomes for Helicobacter pylori Infections. Ann Intern Med 139(6):463.

22. Catry B, et al. (2018) Characteristics of the antibiotic regimen that affect antimicrobial resistance in urinary pathogens. Antimicrob Resist Infect Control 7(1). doi:10.1186/s13756-018-0368-3.

23. Huttner B, Samore M (2011) Outpatient Antibiotic Use in the United States: Time to “Get Smarter.” Clin Infect Dis 53(7):640–643.

24. Schechner V, Temkin E, Harbarth S, Carmeli Y, Schwaber MJ (2013) Epidemiological interpretation of studies examining the effect of antibiotic usage on resistance. Clin Microbiol Rev 26(2):289–307.

25. Truven Health MarketScan Database (2015) Commercial Claims and Encounters (Ann Arbor, MI).

26. Public Health Agency of Sweden, National Veterinary Institute (2015) Consumption of antibiotics and occurrence of antibiotic resistance in Sweden Available at: https://www.folkhalsomyndigheten.se/pagefiles/20281/Swedres-Svarm-2014-14027.pdf.

27. Public Health England (2014) English Surveillance Programme for Antimicrobial Utilisation and Resistance (ESPAUR) Available at: https://www.gov.uk/government/uploads/system/uploads/attachment_data/file/362374/ESP_AUR_Report_2014__3_.pdf.

28. Centers for Disease Control and Prevention (2017) Measuring Outpatient Antibiotic Prescribing. Available at: https://www.cdc.gov/antibiotic-use/community/programs-measurement/measuring-antibiotic-prescribing.html [Accessed February 7, 2018].

29. Hicks LA, et al. (2015) US outpatient antibiotic prescribing variation according to geography, patient population, and provider specialty in 2011. Clin Infect Dis Off Publ Infect Dis Soc Am 60(9):1308–1316.

30. Suskind AM, et al. (2016) Incidence and Management of Uncomplicated Recurrent Urinary Tract Infections in a National Sample of Women in the United States. Urology 90:50–5.

31. Owusu-Edusei K, Carroll DS, Gift TL (2015) Examining Fluoroquinolone Claims Among Gonorrhea-Associated Prescription Drug Claims, 2000-2010. Am J Prev Med 49:761–764.

32. Arizpe A, Reveles KR, Aitken SL (2016) Regional variation in antibiotic prescribing among medicare part D enrollees, 2013. BMC Infect Dis 16(1):744.

33. Bruyndonckx R, et al. (2015) Exploring the association between resistance and outpatient antibiotic use expressed as DDDs or packages. J Antimicrob Chemother 70(4):1241–1244.

34. Hicks LA, Taylor TH, Hunkler RJ (2013) U.S. outpatient antibiotic prescribing, 2010. N Engl J Med 368:1461–2.

35. MacFadden DR, et al. (2016) A Platform for Monitoring Regional Antimicrobial Resistance, Using Online Data Sources: ResistanceOpen. J Infect Dis 214(suppl_4):S393–S398.

36. Wang A, Daneman N, Tan C, Brownstein JS, MacFadden DR (2017) Evaluating the Relationship Between Hospital Antibiotic Use and Antibiotic Resistance in Common Nosocomial Pathogens. Infect Control Hosp Epidemiol 38:1457–1463.

37. Hindler JF, Stelling J (2007) Analysis and presentation of cumulative antibiograms: a new consensus guideline from the Clinical and Laboratory Standards Institute. Clin Infect Dis 44:867–73.

38. MacDougall C, Powell JP, Johnson CK, Edmond MB, Polk RE (2005) Hospital and community fluoroquinolone use and resistance in Staphylococcus aureus and Escherichia coli in 17 US hospitals. Clin Infect Dis 41:435–40.

39. Kanjilal S, et al. (2017) Trends in antibiotic susceptibility in Staphylococcus aureus in Boston, Massachusetts, 2000-2014. J Clin Microbiol. doi:10.1128/JCM.01160-17.

40. Garcia-Rodriguez JA, Fresnadillo Martinez MJ (2002) Dynamics of nasopharyngeal colonization by potential respiratory pathogens. J Antimicrob Chemother 50 Suppl S2:59–73.

41. Dartmouth Atlas of Health Care Research methods.

42. Schisterman EF, Perkins NJ, Mumford SL, Ahrens KA, Mitchell EM (2017) Collinearity and Causal Diagrams: A Lesson on the Importance of Model Specification. Epidemiol Camb Mass 28(1):47–53.

43. Priest P, Wise R, Yudkin P, McNulty C, Mant D (2001) Antibacterial prescribing and antibacterial resistance in English general practice: cross sectional study. BMJ 323(7320):1037–1041.

44. Magee JT, Pritchard EL, Fitzgerald KA, Dunstan FD, Howard AJ (1999) Antibiotic prescribing and antibiotic resistance in community practice: retrospective study, 1996-8. BMJ 319:1239–40.

45. Livermore DM, et al. (2000) Regional variation in ampicillin and trimethoprim resistance in Escherichia coli in England from 1990 to 1997, in relation to antibacterial prescribing. J Antimicrob Chemother 46(3):411–422.

46. Tedijanto C, Olesen SW, Grad Y, Lipsitch M Estimating the proportion of bystander selection for antibiotic resistance among potentially pathogenic bacterial flora. Proc Natl Acad Sci.

47. Hicks LA, Chien Y-W, Taylor TH, Haber M, Klugman KP (2011) Outpatient Antibiotic Prescribing and Nonsusceptible Streptococcus pneumoniae in the United States, 1996–2003. Clin Infect Dis 53(7):631–639.

48. Knight GM, et al. (2018) Quantifying where human acquisition of antibiotic resistance occurs: a mathematical modelling study. BMC Med 16(1):137.

49. Lipsitch M (2001) The rise and fall of antimicrobial resistance. Trends Microbiol 9(9):438–444.

50. Hennessy TW, et al. (2002) Changes in antibiotic-prescribing practices and carriage of penicillin-resistant Streptococcus pneumoniae: A controlled intervention trial in rural Alaska. Clin Infect Dis 34:1543–50.

51. Pouwels KB, et al. (2018) Association between use of different antibiotics and trimethoprimresistance: going beyond the obvious crude association. J Antimicrob Chemoth. doi:10.1093/jac/dky031.

52. Weber SG, Gold HS, Hooper DC, Karchmer AW, Carmeli Y (2003) Fluoroquinolones and the risk for methicillin-resistant Staphylococcus aureus in hospitalized patients. Emerg Infect Dis 9:1415–22.

53. Van Boeckel TP, et al. (2014) Global antibiotic consumption 2000 to 2010: an analysis of national pharmaceutical sales data. Lancet Infect Dis 14(8):742–750.

54. Steiner JF, Prochazka AV (1997) The assessment of refill compliance using pharmacy records: methods, validity, and applications. J Clin Epidemiol 50:105–116.

55. Harbarth S, Harris AD, Carmeli Y, Samore MH (2001) Parallel analysis of individual and aggregated data on antibiotic exposure and resistance in gram‐begative bacilli. Clin Infect Dis 33:1462–1468.

56. Lipsitch M, Samore MH (2002) Antimicrobial use and antimicrobial resistance: a population perspective. Emerg Infect Dis 8:347–354.

57. B. R. Levin, et al. (1997) The population genetics of antibiotic resistance. Clin Infect Dis 24. doi:10.1093/clinids/24.supplement_1.s9.

58. Colijn C, et al. (2010) What is the mechanism for persistent coexistence of drug-susceptible and drug-resistant strains of Streptococcus pneumoniae? J R Soc Interface 7(47):905–919.

59. Olesen SW, Barnett ML, MacFadden DR, Lipsitch M, Grad YH (2018) Trends in outpatient antibiotic use and prescribing practice among US older adults, 2011-15: observational study. BMJ:k3155.

60. Benjamini Y, Hochberg Y (1995) Controlling the false discovery rate: a practical and powerful approach to multiple testing. J R Stat Soc Ser B Stat Methodol 57:289–300.

